# The Carbon Starvation-Inducible Lipoprotein (Slp) Influences Differential Adherence of *Escherichia coli* O157:H7 at the Bovine Rectoanal Junction

**DOI:** 10.1101/2025.10.02.679962

**Authors:** Indira T. Kudva, Erika N. Biernbaum, Eric D. Cassmann, Mitchell V. Palmer, Lekshmi K. Edison, Jessy Castellanos-Gell, Subhashinie Kariyawasam

## Abstract

Shiga toxin-producing *Escherichia coli* O157:H7 (O157), a foodborne human pathogen, persists at the rectoanal junction (RAJ) of the bovine intestinal tract, in asymptomatic cattle reservoirs. Identifying mechanisms used by O157 for initial adherence before persistence at the RAJ could help develop effective O157 control modalities. We recently established the role of carbon starvation-inducible lipoprotein (Slp) in initial adherence of O157 to Caco-2 cells, with the human polymeric immunoglobulin receptor (pIgR) protein as the Slp-receptor. Here, we evaluated the role of Slp in O157 adherence to the bovine RAJ using the RAJ squamous epithelial (RSE) cell- and RAJ-*In vitro* Organ Culture (IVOC)- adherence assays. The wild-type O157 strain EDL932 (EDL932-WT), it’s isogenic *slp* deletion mutant (EDL932 Δ*slp*), and the *slp* complemented mutant (EDL932 Δ*slp*-p:*slp*), were tested with no bacteria controls. Adherence was verified by culture and immunofluorescence (IF) staining of O157. Tissue integrity was determined using nuclear/cell staining dyes and histopathological examination. All test strains adhered in a diffuse-moderate pattern on RSE cells. However, differential adherence was observed on the RAJ-IVOC with the strains preferentially adhering to the columnar cells. Additionally, EDL932- WT and EDL932 Δ*slp*-p:*slp* strains adhered in slightly greater numbers than the EDL932 Δ*slp* strain to the RAJ-IVOC, causing disruptions primarily in the columnar region of otherwise intact RAJ-IVOC tissues. Interestingly, pIgR was also predominantly detected by IF microscopy and RNAscope in situ hybridization at the columnar region of the RAJ-IVOC tissue. *In silico* modeling demonstrated the possibility of a bovine pIgR- bacterial Slp interaction. Hence, our observations support the role for Slp in the initial adherence of O157 to the columnar cells at the bovine RAJ, unlike the squamous cells where the loss of *slp* did not impact attachment. In addition, a possible mucosal immune-interference resulting from the bovine pIgR-Slp interaction could contribute towards long-term O157 colonization of cattle.

**AUTHOR SUMMARY:** *Escherichia coli* O157:H7 (O157) is a foodborne pathogen that causes disease in humans with symptoms ranging from watery/bloody diarrhea to kidney failure. Cattle are the main reservoirs of this human pathogen, where O157 tends to persist at the rectoanal junction (RAJ) of the bovine gastrointestinal tract. However, the exact mechanisms by which O157 adheres and persists at the bovine RAJ is not fully understood. Here, we identified that the O157 carbon starvation-inducible lipoprotein (Slp) allows O157 to attach to cells at the RAJ using polymeric immunoglobulin receptor (pIgR) protein as the Slp-receptor. Not only does this attachment of Slp to the cattle receptor allow O157 to colonize cattle, but the interaction could potentially interfere with intestinal immune responses, further promoting long-term O157 colonization of cattle. This makes Slp an attractive target for therapeutic interventions, such as vaccines that could interfere with O157 attachment to cattle intestinal cells.

## INTRODUCTION

Shiga toxin-producing *Escherichia coli* (STEC) O157:H7 (O157) is a foodborne human pathogen that colonizes the gastrointestinal tracts of cattle, the primary reservoir, without causing disease in the animals [1–3]. However, when humans consume STEC O157-contaminated food, particularly undercooked meat, they can develop severe illnesses ranging from watery to bloody diarrhea that can progress to secondary sequelae such as, hemolytic-uremic syndrome, and even kidney failure [4–8]. In cattle, the preferred site for STEC O157 persistence in the bovine gastrointestinal tract is the rectoanal junction (RAJ) located at the terminal end of the distal colon where the columnar epithelial cells transition to the stratified squamous epithelial cells towards the anus [9–12]. STEC O157 form micro- colonies in a region 3-5 centimeters proximal to the RAJ comprising dense lymphoid follicles covered with columnar epithelial cells referred to as the follicle-associated epithelium (FAE) [9, 13, 14]. In contrast, STEC O157 diffusely adheres to the stratified squamous epithelial cells distal to the RAJ, denoted as the rectoanal junction squamous epithelium (RSE) [15]. Our studies have shown that STEC employ different mechanisms compared to commensal bacteria for attachment to the RAJ; in addition, STEC O157 appear to utilize distinct adherence proteins at the FAE versus RSE regions, which in turn varies from those utilized by the non-O157 STEC [15–20]. Hence, unravelling the mechanisms of STEC O157 interactions with the bovine intestinal cells is crucial for developing optimal strategies that could interfere with cattle colonization, and thereby, minimize human infections.

In STEC O157, as in other *E. coli*, one of the proteins expressed during nutrient limitation is a carbon starvation and stationary-phase inducible lipoprotein, Slp. This is a 22 kDa lipoprotein associated with the *E. coli* outer membrane [21–23]. The *slp* gene is located on a genomic acid fitness island (AFI) in all *E. coli*, although in STEC the AFI has an insertion altering its size and possibly regulation [24–26]. The primary role of Slp is to stabilize the bacterial outer membrane under nutrient limiting conditions [21, 23], although it is also associated with improved uptake of nutrients during starvation [22], biofilm formation [27], protection against hydrogen peroxide stress [28], and as a putative receptor for the lambda phage protein, NinD [29]. Interestingly, in our previous study, Slp was identified as one of the proteins expressed by STEC O157 in Dulbecco’s modified Eagle’s medium, a nutrient limited media, and was predicted to be a putative adhesin using Vaxign, a reverse vaccinology-based vaccine target prediction and analysis system (http://www.violinet.org) [16]. Subsequently, in another study, we demonstrated that Slp plays a role in the initial adherence of STEC O157 to the human colonic epithelial cells [26]; when the *slp* gene was disrupted, it led to a significant reduction in initial adherence to Caco-2 cells, a type of human colonic cell line. This effect was reversed by adding a plasmid containing the *slp* gene, and overexpression of Slp even lead to increased adherence compared to the wild-type STEC O157 [26].

In the same study, the polymeric immunoglobulin receptor was determined to be a receptor for Slp on Caco2-cells using co-localization experiments [26]. The polymeric immunoglobulin receptor (pIgR) is an eukaryotic glycoprotein that can vary in size from 80 to 120 kDa, based on the level of glycosylation, and is involved in the transportation of polymeric immunoglobulin A (IgA) and IgM from the basolateral to the apical surface of mucosal epithelial cells via transcytosis, contributing to gut health and homeostasis [30–34]. In humans, pIgR is localized primarily in the intestinal tissue with a predominance in the duodenum and colon [35] and in few instances, antibody-free, membrane-bound pIgR on the apical surface of mucosal epithelia can get recycled via a retrograde pathway [30–32, 34–36]. This retrograde uptake of pIgR is exploited by pathogens such as *Streptococcus pneumoniae* that invade the nasopharyngeal cells by binding pIgR with the PspC adhesin [36, 37].

Our observation of the attachment of STEC O157 to pIgR via the Slp lipoprotein was unique and not observed with *E. coli* K12, highlighting a potential mechanism by which STEC O157 can utilize starvation-induced adaptations to establish initial colonization in the host [26]. Since the Caco2 cells are of colonic derivatization, we hypothesized that similar Slp-pIgR interactions could be occurring at the bovine RAJ at end of the distal colon in cattle. Li et al in their studies evaluating genes expressed at the bovine RAJ following STEC challenge of cattle observed an upregulation of pIgR at 6h post-challenge, especially with non-O157 STEC, suggesting presence of a local mucosal immune response at the lymphoid follicular region of the RAJ [38]. In our comparative transcriptome studies, we observed an upregulation of Slp in STEC O157 adhering *in vitro* to both human colonic (CCD CoN 841) and RAJ epithelial cells [39]. Since the pH at the distal colon-RAJ is close to neutral, it is likely that nutrient limitation could be the major factor contributing to the upregulation of Slp expression in STEC O157, and the fecal-RAJ microbiota stimulating overall pIgR expression by the mucosal epithelia at this site [8, 21, 23, 30, 40, 41, 42] making it conducive for Slp-pIgR interactions. In this study, we ascertained if Slp would indeed play a role in the initial attachment of STEC O157 to the bovine RAJ using the RSE cells and RAJ-*In vitro* Organ Culture (IVOC) *in vitro* adherence assays and verified the possible role of pIgR expression in this adherence.

## RESULTS

### Differential adherence patterns were observed on the RAJ-IVOC at the 10^7^ CFU inoculum unlike with RSE cells only

All three test strains, inoculated at a bacteria:cell ratio of 10:1, demonstrated the same diffuse, moderate adherence pattern on RSE cells with no significant quantitative differences between strains (70 – 76% cells having 1-10 bacteria/cell; *p=*0.8 -0.7), irrespective of the presence or absence of *slp* (Fig. 1). This pattern was observed on the squamous epithelial cells of the RAJ-IVOC tissue as well, however, a distinct increased adherence to the columnar region of the same tissue was observed that appeared to be linked to the presence/absence of *slp* (Figs. 2 and 3).

**Figure 1.**
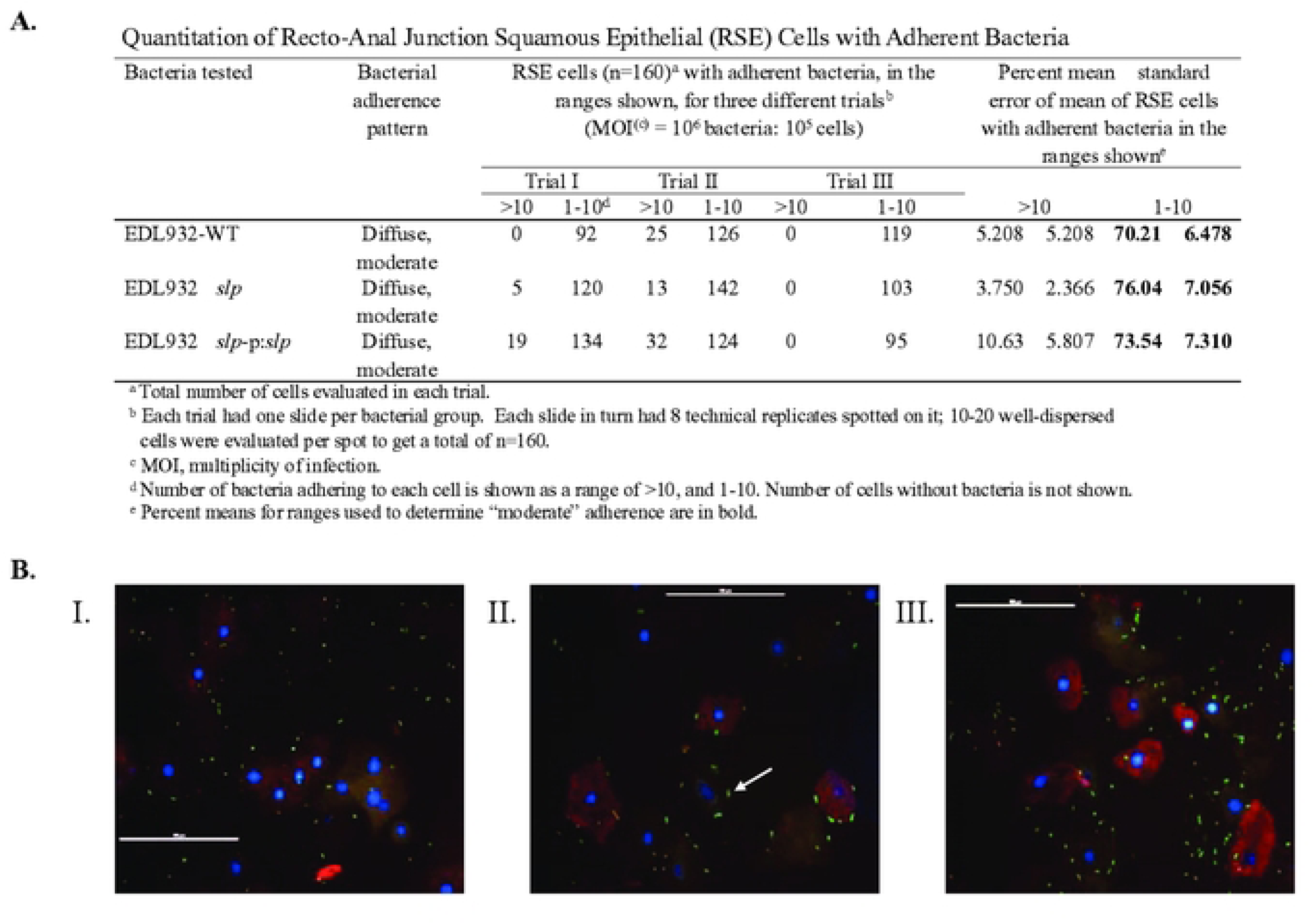
**Quantitative (Panel A) and qualitative (Panel B) data from the RSE cell adherence assay.** The quantitative data from three adherence assays are shown in the table in Panel A. The ‘diffuse, moderate’ adherence patterns of the test strains on the RSE cells are shown in Panel B. The immunofluorescent images were captured at 400x magnification with the 100 µm scale bar. The bacteria (STEC O157) as indicated by arrows, the RSE cells’ cytokeratins, and the nuclei have green, orange-red and blue fluorescence, respectively.

**Figure 2.**
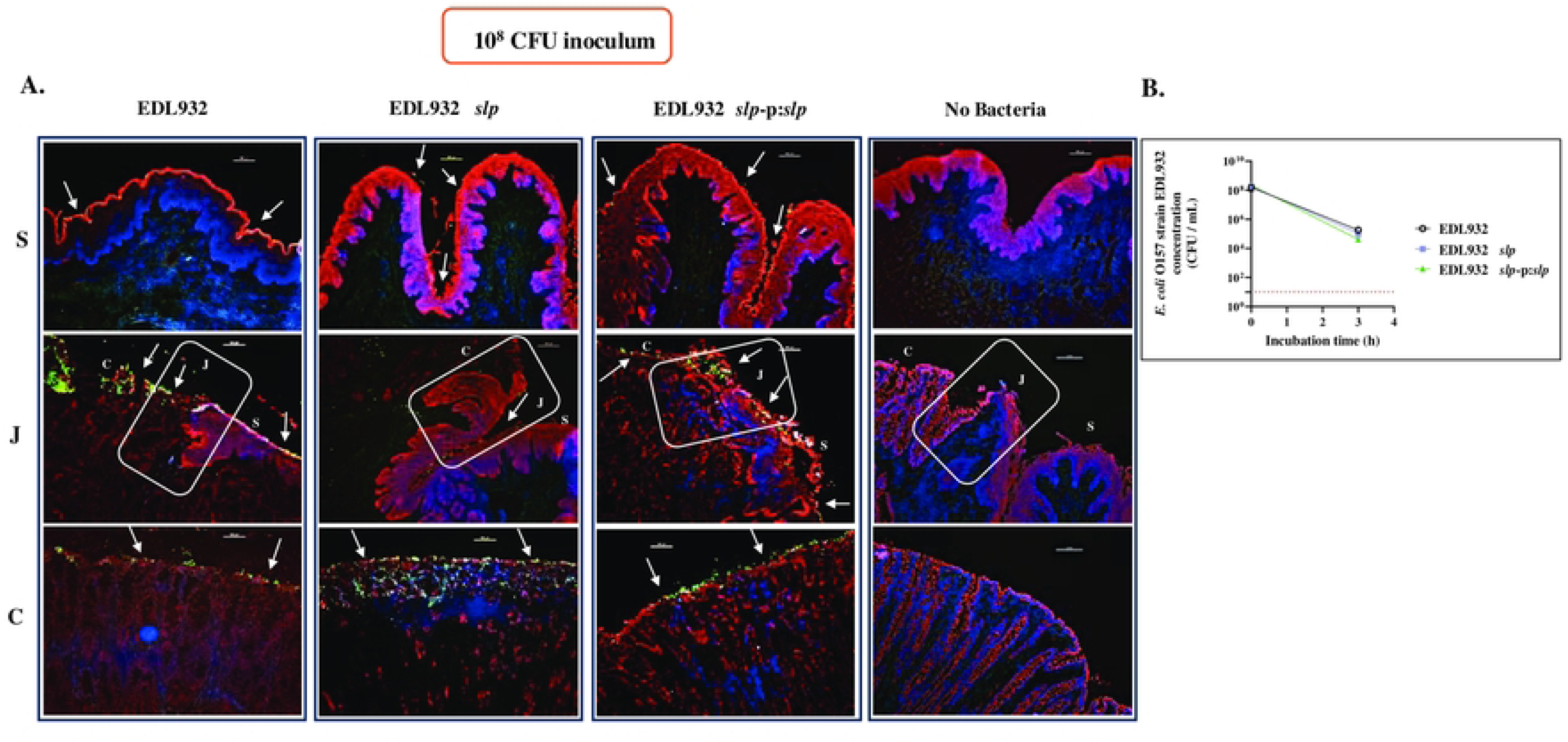
Immunofluorescent RAJ-IVOC images and graph depicting STEC O157 adherence and viable counts of recovered bacteria, respectively, post inoculation with 10^8^ CFU test strains. **(A)** Immunofluorescent images of tissue sections from RAJ-IVOCs used to set up the adherence assay with 10^8^ CFU inoculum concentration for all the test strains or not inoculated (no bacteria). Post-assay, tissue sections were stained with immunofluorescent antibodies targeting the RAJ cells’ cytokeratin and STEC O157, and images were recorded at 100x magnification. The adherent bacteria (shown with arrows), RAJ cells’ cytokeratin, and the nuclei have green, orange-red and blue fluorescence, respectively. The squamous (S), junction (J) and columnar (C) regions of the RAJ are indicated, along with a 100 µm scale bar. **(B)** Cumulative graph from two assays, showing viable counts (CFU/ml) of test strains recovered from RAJ-IVOC tissues by bacterial culture. The red dotted line on the graph marks the STEC O157 detection limit of 10 CFU/ml for non-enrichment cultures.

**Figure 3.**
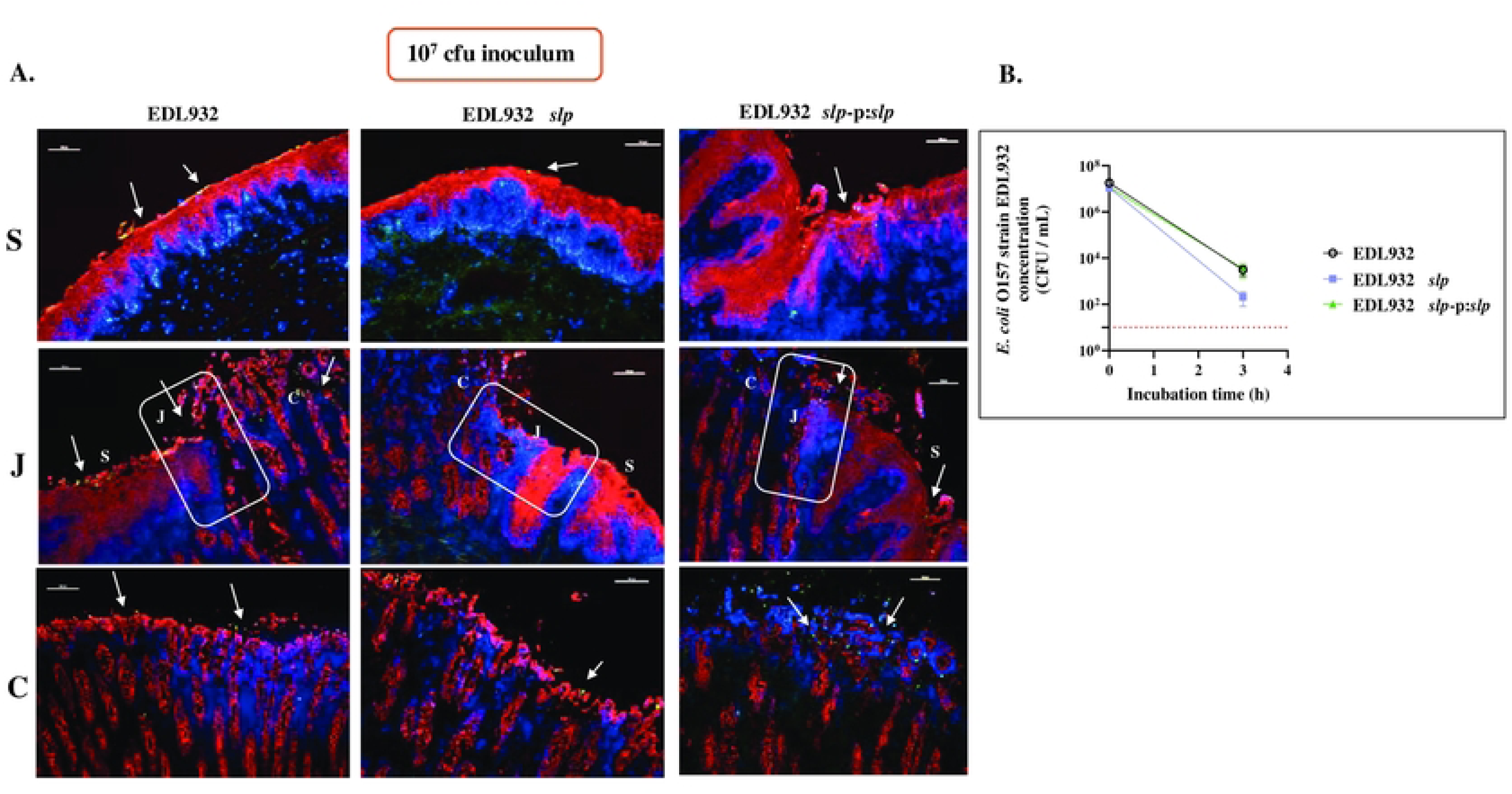
Immunofluorescent RAJ-IVOC images and graph depicting STEC O157 adherence and viable counts of recovered bacteria, respectively, post inoculation with 10^7^ CFU test strains. **(A)** Immunofluorescent images of tissue sections from RAJ-IVOCs used to set up the adherence assay with 10^7^ CFU inoculum concentration for all the test strains. Post-assay, tissue sections were stained with immunofluorescent antibodies targeting the RAJ cells’ cytokeratin and STEC O157, and images were recorded at 100x magnification. The adherent bacteria (shown with arrows), RAJ cells’ cytokeratin, and the nuclei have green, orange-red and blue fluorescence, respectively. The squamous (S), junction (J) and columnar (C) regions of the RAJ are indicated, along with a 100 µm scale bar. **(B)** Cumulative graph from four assays showing viable counts (CFU/ml) of test strains recovered from RAJ-IVOC tissues by bacterial culture. The red dotted line on the graph marks the STEC O157 detection limit of 10 CFU/ml for non-enrichment cultures.

For the RAJ-IVOC, two different concentrations of the test strains were used in the inoculum, 10^8^ and 10^7^ CFU in 2ml media, to determine the optimal inoculation dose (Figs. 2 and 3). The lower inoculum concentration allowed for better clarity in bacterial adherence phenotype and hence, the 10^7^ CFU in 2 ml media inoculum was used in all subsequent six RAJ-IVOC assays. Immunofluorescent imaging of tissue sections of the frozen RAJ-IVOCs indicated the differential adherence pattern of the test strains, with EDL932-WT and EDL932 Δ*slp*-p:*slp* adhering in greater numbers along the columnar epithelia especially along the FAE cells compared to EDL932 Δ*slp* (Figs 2 and 3). The adherence phenotype was primarily diffuse along the squamous cells but diffuse with several aggregates on the columnar epithelia of the RAJ-IVOC (Figs 2 and 3). Increased focal disruption of epithelia, especially around the junction and the columnar epithelia, could be associated only with the presence of STEC O157 (Figs. 2 and 3; Supp. Fig. S5 and Supp. Histopathology Reports).

Post-incubation, culture of both the leftover inoculum and the tissue itself enabled the recovery of the test strains from the respective RAJ-IVOCs. The viable counts from the residual inoculum ranged from 10^3^ to 10^7^ CFU/ml for each test strain (Supp. Table S1). On the other hand, there was a reduction in the viable counts for the test strains recovered from the tissues, suggesting loss of non-adhered bacteria to the washes prior to processing tissues for culture. On an average, about 2- to 3- log decrease in viable counts was observed with the EDL932-WT and EDL932 Δ*slp*-p:*slp* strains, and a 4- to 5- log reduction in the EDL932 Δ*slp* viable counts (Fig. 3; Supp. Table S1 and Fig. S2) when the 10^7^ CFU inoculum was used. This could reflect the poor adherence by the EDL932 Δ*slp* strain, as observed in the immunofluorescent images, although the differences in counts were not statistically significant *(p* = 0.5 - 0.9). These differences in the recovery of the tissue adherent-bacteria by culture were not discernable with the 10^8^ CFU inoculum where a reduction by 2- to 3- logs was observed in all viable counts irrespective of the strain (Fig. 2; Supp. Table S1 and Fig. S2).

No STEC O157 were detected in the no bacteria controls set up for the RSE cell- and RAJ-IVOC -adherence assays eliminating the possibility of any pre-existing or cross-contaminating STEC O157. Occasionally, a few pinpoint background colonies were obtained on culture plates that were ruled out to be STEC or *E. coli* based on the phenotype and serological testing. All recovered bacteria produced sorbitol non-fermenting, colorless, MUG-non-utilizing, non-fluorescent colonies typical of STEC O157 and readily agglutinated with the O157 latex agglutination reagent. Additional PCR verified the genotype of the recovered test strains. The distinctive polymorphic amplified typing sequence (PATS) profiles for the STEC O157 strain EDL932 helped confirm the relatedness of the wild type, mutant, and complemented strains used in this study (Table 1A; Supp. Fig. S3). In addition, the PCR results with the vector and *slp* -gene primers distinguished the EDL932 Δ*slp* and EDL932 Δ*slp*-p:*slp* strains from the parent EDL932-WT (Table 1B and Supp. Fig. S3).

**Table 1A.** Representative PATS profiles of isolates inoculated and recovered from RAJ-IVOCs.

| Strain | Polymorphic *Xba*I sites |  |  |  |  |  |  |  | Polymorphic *Avr*II sites |  |  |  |  |  |  | Virulence Genes |  |  |  |
| --- | --- | --- | --- | --- | --- | --- | --- | --- | --- | --- | --- | --- | --- | --- | --- | --- | --- | --- | --- |
|  | IK8 | IK25 | IK114 | IK118 | IK123 | IK127 | IKB3 | IKB5 | IKNR3 | IKNR7 | IKNR10 | IKNR12 | IKNR16 | IKNR27 | IKNR33 | *stx1* | *stx2* | *eaeA* | *hlyA* |
| EDL932-WT | 0 | 0 | 1 | 1 | 1 | 1 | 1 | 1 | 2 | 2 | 2 | 2 | 2 | 2 | 2 | 1 | 1 | 1 | 1 |
| EDL932 Δ*slp* | 0 | 0 | 1 | 1 | 1 | 1 | 1 | 1 | 2 | 2 | 2 | 2 | 2 | 2 | 2 | 1 | 1 | 1 | 1 |
| EDL932 Δ*slp*-p:*slp* | 0 | 0 | 1 | 1 | 1 | 1 | 1 | 1 | 2 | 2 | 2 | 2 | 2 | 2 | 2 | 1 | 1 | 1 | 1 |

**Table 1B.** Representative PCR profiles of isolates recovered from RAJ-IVOCs: Post assay, in leftover inoculum (post) and from non- enrichment bacterial culture of tissues (TC-NE).

| Isolate | *slp* | pUC18 | *stx2* | Match with the corresponding test strain |
| --- | --- | --- | --- | --- |
| EDL932-WT-A post | + | - | + | ✓ |
| EDL932-WT-B post | + | - | + | ✓ |
| EDL932 Δ*slp*-A post | - | - | + | ✓ |
| EDL932 Δ*slp*-B post | - | - | + | ✓ |
| EDL932 Δ*slp*-p:*slp*-A post | + | + | + | ✓ |
| EDL932 Δ*slp*-p:*slp*-B post | + | + | + | ✓ |
| EDL932-WT-A TC-NE | + | - | + | ✓ |
| EDL932-WT-B TC-NE | + | - | + | ✓ |
| EDL932 Δ*slp*-A TC-NE | - | - | + | ✓ |
| EDL932 Δ*slp*-B TC-SE | - | - | + | ✓ |
| EDL932Δ*slp*-p:*slp*-A TC-SE | + | + | + | ✓ |
| EDL932Δ*slp*-p:*slp*-B TC-NE | + | + | + | ✓ |

### The RAJ-IVOC tissue integrity was disrupted primarily by the STEC O157 inoculum

The RedDot2 nuclear stain confirmed the viability of cells within the RAJ-IVOC with the absence of nuclear staining in the un-fixed RAJ-IVOC tissues pre- and post- 3 h incubation (Supp. Fig. S4). Additional histopathological evaluations linked epithelial disruption in inoculated IVOCs primarily to the presence of STEC O157, as previously reported [43]. The squamous and columnar epithelial cell regions along with the junction were clearly visible in all H&E-stained RAJ-IVOC tissue sections along with lymphoid follicles underlying some of the columnar epithelia (FAE cells); the latter region being referred to as the glandular or mucosal region in some of the pathology reports (Supp. Histopathology Reports folder).

Some animals had antemortem anomalies and/or disease contributing to histopathology in parts of the RAJ limiting extensive use of the tissue in setting up IVOCs (Supp. Histopathology Reports folder). However, as shown in the representative histopathological report (Supp. Fig. S5), overall, the glandular and squamous regions were normal/unremarkable in the tissue sections used to set up the no bacteria- control RAJ-IVOC (Supp. Fig. S5). Additional epithelial disruptions were observed largely in the glandular region of RAJ-IVOC exposed to EDL932-WT, EDL932 Δ*slp*, or EDL932 Δ*slp*-p:*slp*, respectively (Supp. Fig. S5; Supp. Histopathology Reports folder).

### pIgR localizes primarily along the columnar epithelial region of the RAJ-IVOC with likely suppression after Slp binding

Immunofluorescent staining of the RAJ-IVOC tissue sections for pIgR indicated a predominant distribution of this receptor in the columnar epithelial region of the RAJ with minimal-to-no pIgR along the squamous epithelia (Fig. 4). pIgR could be detected both within and on the luminal surface of the columnar epithelia (Fig. 4). Hence, the increased adherence of *slp*-expressing EDL932-WT and EDL932 Δ*slp*-p:*slp* to the columnar epithelia (Figs. 2 and 3) appeared to also correlate with the increased expression of host cell pIgR (Fig. 4) in that region. This biased distribution of pIgR on the RAJ-IVOC tissues was also observed by RNAScope in situ hybridization (ISH); distinct spatial expression of the pIgR mRNA was detected primarily in the cytoplasm of columnar cells in all the RAJ- IVOC tissue sections evaluated (Supp. Table S2 and Supp. RNAScope Images folder).

**Figure 4.**
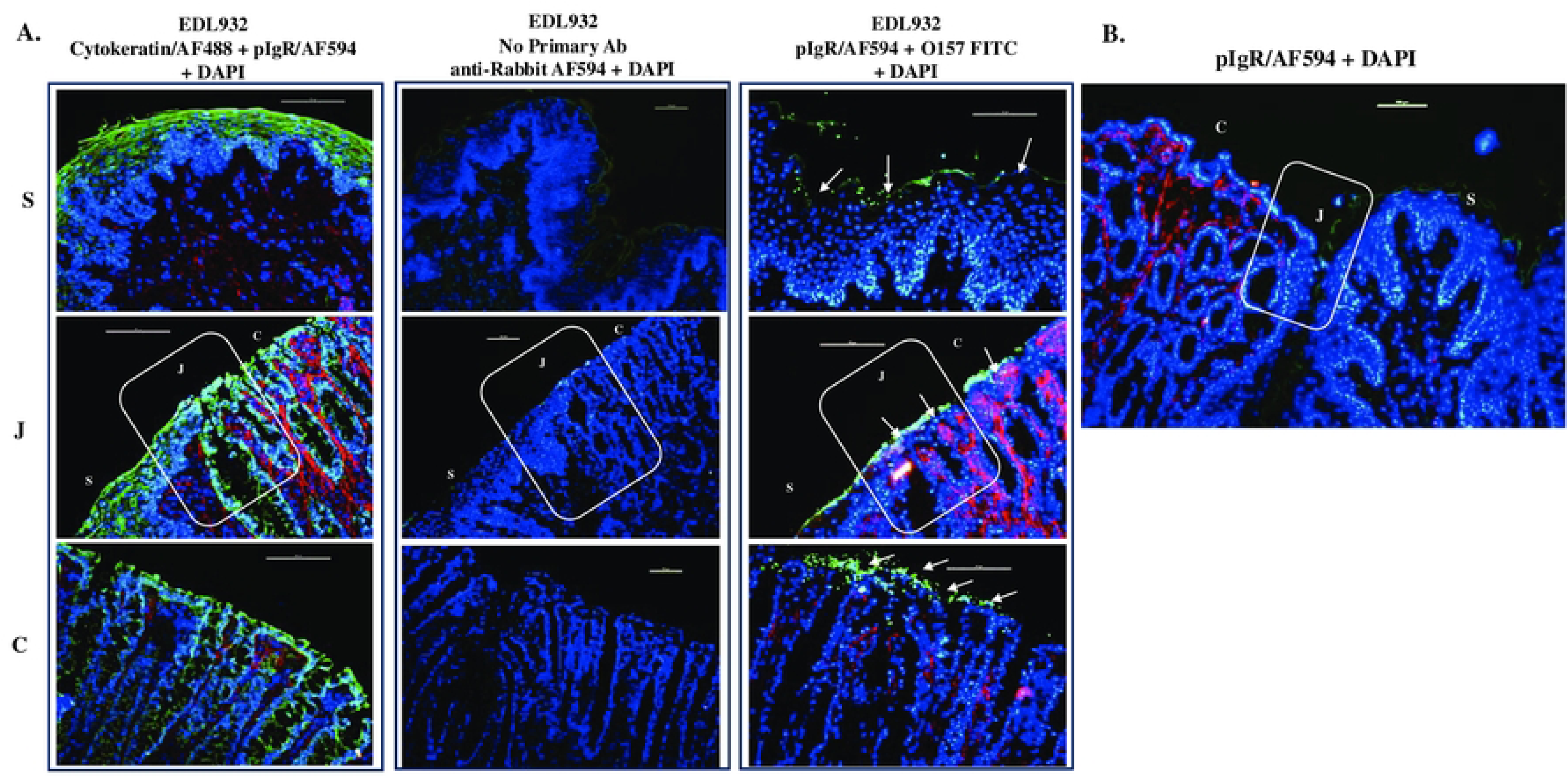
Representative immunofluorescent images of tissue sections from the RAJ-IVOC adherence assay depicting pIgR distribution. **(A)** RAJ-IVOC were inoculated with EDL932-WT (10^7^ CFU inoculum) and post-assay tissue sections were then stained with immunofluorescent antibodies targeting pIgR, STEC O157/or RAJ cells’ cytokeratins, and images recorded at 200x magnification. The pIgR (red), adherent bacteria (green; shown with arrows) or RAJ cells’ cytokeratins (green), and the nuclei (blue) are shown. The squamous (S), junction (J) and columnar (C) regions of the RAJ are indicated, along with a 100 µm scale bar. A ‘no primary antibody’ control was included to demonstrate the specificity of the antibodies targeting pIgR**. (B)** The additional image depicting the regions around the junction demonstrates the predominant distribution of pIgR in the columnar epithelium of the RAJ tissue.

In all the RNAScope ISH assays, the positive control probe targeting the cyclophilin mRNA generated robust signals verifying RNA integrity and the negative control dapB mRNA-targeting probe generated no signal indicating lack of any cross-reacting RNA in the background (Supp. RNAScope Images folder). In the trial run, an unexpected low ‘average red (pIgR) copies per cell’ was obtained for the single columnar epithelial (SCE) region of the RAJ-IVOC tissue exposed to EDL932 WT (**#3**: 0.538965) compared to the higher scores for tissues exposed to EDL932 Δ*slp* **(#4:** 4.946465) or not exposed to any inoculum (Pre-assay control **T0:** 1.281183 and no bacteria control **#1:** 2.491031) (Supp. Table S2-Trial Run). Similar output was obtained in the test run with a different set of RAJ-IVOC tissues. For the test run, the ‘average red (pIgR) copies per cell’ scores for the SCE regions were: **T0:** 3.409, **#1:** 2.374, **#3:** 1.760, **#4:** 4.169, **#5:** 1.853 (Supp. Table S2-Test Results). Such distinct differences were not observed in the stratified squamous epithelial (SSE) regions. Overall, these results provide additional evidence of the RAJ-IVOC tissue viability and most importantly suggest a possible suppression of pIgR in presence of the interacting ligand, Slp. Further studies are needed to explore this observation as it may provide insights into the reduced mucosal (IgA) responses in STEC colonized or STEC-vaccinated cattle [33, 44, 45].

### Protein-protein docking predictions support pIgR-Slp interaction

A total of 4 and 10 possible 3D-docking models were generated in NovaDock, depicting the human pIgR-*E. coli* Slp and bovine pIgR- *E.coli* Slp interactions, respectively. Model energy, cluster size and cluster energy was considered when selecting an optimal model (Fig. 5; Supp. PDB files folder); a low energy and high cluster size is usually indicative of low energy and biologically relevant conformation (https://www.dnastar.com; [46–48]).

**Figure 5.**
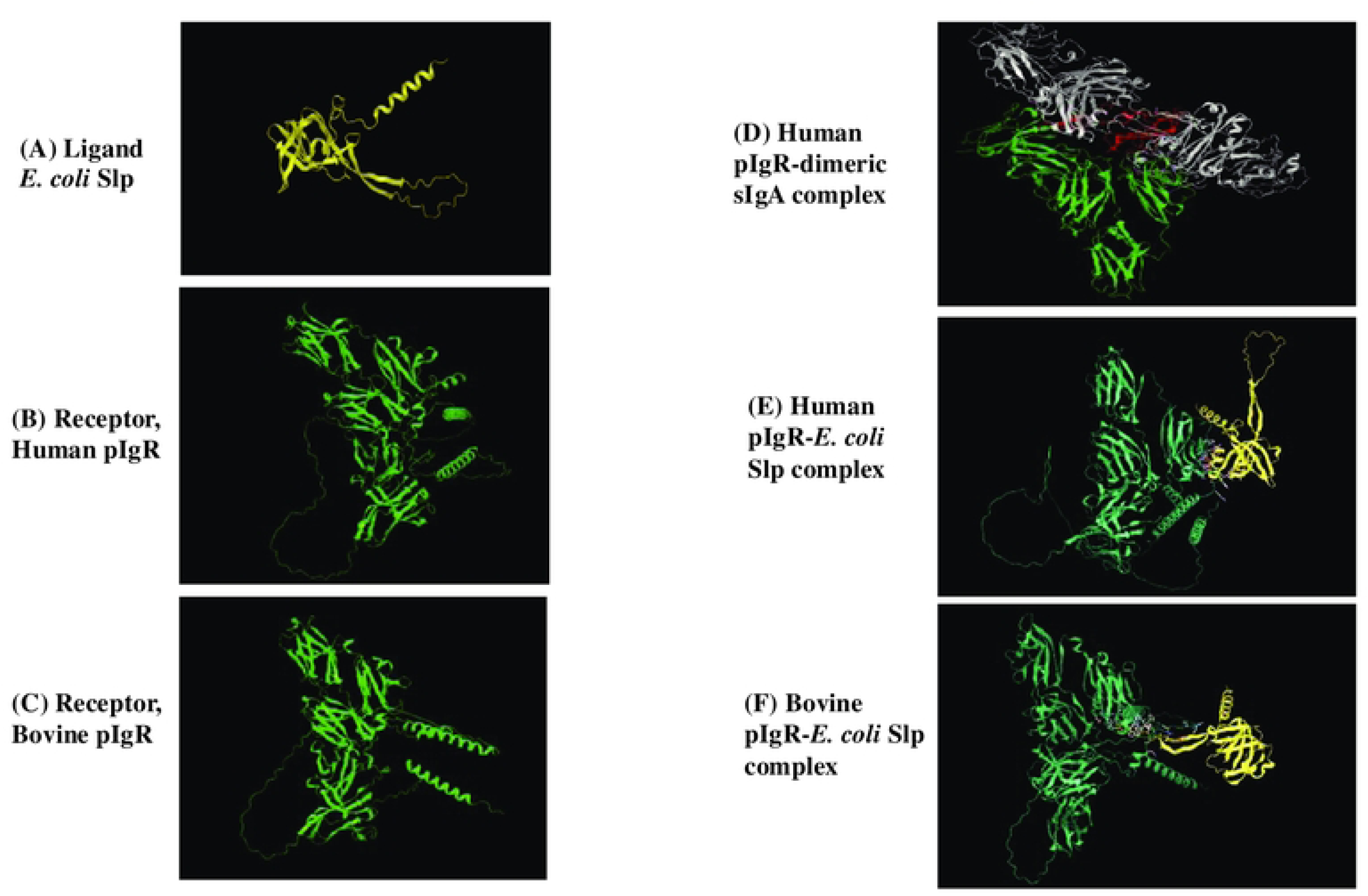
Representative computational models for individual and interacting proteins. (A) Ligand *E. coli* Slp. (AlphaFold), **(B) Receptor, Human pIgR** (AlphaFold), **(C) Receptor, Bovine pIGR** (AlphaFold), **(D) Human pIgR-dimeric sIgA complex** (RCSB), **(E) NovaDock predicted model: Human pIgR-*E. coli* Slp complex**, and **(F) NovaDock predicted model: Bovine pIgR-*E. coli* Slp complex**. Color style used was: Yellow-Slp; Green-pIgR; White-dimeric sIgA; Red-J chain.

Details of the selected models for each of the predicted receptor-ligand combinations are in the supplementary document with the additional model information, providing a list of residues involved in intermolecular contacts at the binding interface (Supp. Additional model and BLASTp information). The parameters of the best scoring docking model for the human pIgR-*E. coli* Slp was, model energy: -18.30, cluster size: 2, cluster energy: -12.01± 6.29 with 40 residue contacts (Supp. Additional model and BLASTp information). Likewise, the parameters for the optimal bovine pIgR-*E. coli* Slp docking model was, model energy: -34.95, cluster size: 1, cluster energy: -34.951± 0 with 41 residue contacts (Supp. Model Information). Although the cluster size was limited, the low energy of the conformations support optimal interactions between ligand and receptor. Interestingly, BLASTp analysis indicated a homology of only 67% between the human and bovine pIgR sequences (Supp. Additional model and BLASTp information), which may have contributed to differences in the Slp-docking models and interacting residues. Overall, as expected, fewer residues were involved in pIgR-Slp docking compared to the more traditional pIgR- dimeric sIgA complex (Fig. 5; Supp. PDB files folder).

## DISCUSSION

STEC O157 are armed with several fimbrial and non-fimbrial adhesins that are either unique to STEC or shared with other *E. coli* [8, 49–57]. While most of these adhesins are associated with transient adherence, the STEC adhesin intimin-γ enables intimate attachment to the colonic epithelia leading to the onset of hemorrhagic colitis in humans through cellular damage [5, 54, 58–67]. Intimin-γ, has also been associated with the tissue-tropism of STEC to the FAE cells of the RAJ in cattle [9, 53]. In addition, *in vitro* evaluation of STEC O157 in bovine digestive contents from rumen, small intestine and rectum demonstrated niche-specific expression of intimin-γ, and other adhesins [68, 69]. However, our research excluded the role of intimin-γ in STEC adherence to the RSE cells at the RAJ suggesting a role for alternate adhesins [16, 17, 70, 71]. Our investigations revealed a variable (host-, host cell type- or bacterial strain -specific), and/or a modulating role for some of the routinely studied adhesins including Intimin-γ, EspA, Curli, YfaL, FimH, Cah autotransporter, OmpA and non-fimbrial adhesins, on RSE cell-adherence [18–20, 26, 39, 57, 70–73].

Hence, in this study, we expanded our investigations to evaluate another non-fimbrial, outer membrane lipoprotein with a role in biofilm formation and adherence to Caco2 cell line, Slp[26, 27]. We included the recently standardized RAJ-IVOC model system, in addition to RSE cells, to ascertain the role of Slp in STEC O157 adherence to all cell-types at the RAJ, namely the squamous and columnar epithelial cells [43]. Interestingly, we observed a slightly increased adherence by STEC O157 strains expressing Slp, EDL932-WT and EDL932 Δ*slp*-p:*slp*, versus the deletion-mutant, EDL932 Δ*slp*, on the RAJ-IVOC (Fig. 3). This observation was made both by bacterial culture and microscopic examination of the RAJ-IVOCs, although the differences were more discernable by the latter (Fig. 3). A biased, diffuse and aggregative attachment of the tested strains to the columnar epithelial cells was observed while the same strains adhered diffusely to the squamous epithelial cells validating the results of the RSE cell adherence assay (Figs 1, 2 and 3; Table1). The increase or decrease in the adherent bacteria, depending on the strain used, was observed uniformly along the entire RAJ-IVOC tissue (Figs 2 and 3). These results indicate a role for Slp in STEC O157 adherence to the RAJ at the initial stages, as the assays did not extend beyond 3-4 hours of inoculum exposure. Our previous studies had identified pIgR as an Slp- receptor allowing for STEC O157 attachment to Caco2 cell [26]. This appeared to be congruous with observations made by other researchers of the exploitation of pIgR by respiratory pathogen, *S. pneumoniae* [36, 37]. In the current study, we observed an increased distribution of pIgR, by immunofluorescent microscopy detecting the protein and RNAScope ISH identifying the spatial distribution of the mRNA, along the columnar epithelia where the test strains adhered the most (Figs 2, 3, 5; Supp. Table 2 and Supp. RNAScope Images folder). This prompted computational evaluation using docking-models to determine the viability of the Slp-pIgR interaction. The low energy of the conformations generated for the Slp-pIgR model, supporting optimal interactions between the ligand and receptor, provided biological relevancy.

pIgR expression is usually stimulated by the membrane lipopolysaccharides of commensal Gram-negative bacteria through the toll-like receptor 4 (TLR4) for reduced inflammatory responses that can be detrimental to bacteria and host to some extent [30, 40]. Li et al observed a more pronounced upregulation of pIgR at the RAJ of 5–10-month-old Holstein steers challenged with non-O157 STEC, irrespective of the colonizing ability, compared to the colonizing STEC O157 [38]. We used the RAJ from an Aberdeen and a Holstein steer, 1.5 years of age, for the RAJ-IVOC assays, and in both instances, observed a decrease in the pIgR-mRNA in tissues exposed to EDL932-WT or EDL932 Δ*slp*-p:*slp* compared to EDL932 Δ*slp* or the “no bacteria”/ “pre-assay” controls (Supp. Table 2 and Supp. RNAScope Images folder). This differential pIgR expression is similar to the observations made by Li et al with live animals [38], and supports a role for non-O157 STEC/non-STEC Gram-negative bacteria in the stimulation of pIgR expression along with non-Slp expressing STEC O157, which is then exploited by the Slp expressing STEC O157 for initial adherence at the RAJ.

Several plausible reasons support the STEC O157 Slp-host cell pIgR interaction including niche selection, immune evasion and persistence. Binding of Slp to pIgR could contribute towards niche selection by STEC O157 to the columnar epithelium of the RAJ, or similar to *S. penumoniae* this interaction could promote internalization [36, 37] and hence, immune evasion by STEC O157 [74] leading to long term intermittent- or super- shedding by colonized cattle [75–78]. Immune suppression and STEC persistence at the RAJ with increased shedding have previously been linked [44, 45, 79–81]. Slp binding of pIgR could suppress pIgR-mRNA expression as observed with the enterotoxigenic *Escherichia coli* in a mouse disease model by Liu et al [82] as an immune-evasion tactic [33] and may explain the decrease in pIgR-mRNA in tissues exposed to EDL932-WT or EDL932 Δ*slp*-p:*slp* compared to EDL932 Δ*slp* or the controls with no inoculum (Supp. Table 2 and Supp. RNAScope Images folder). Slp binding of some ‘free’ pIgR may also interfere with the optimal apical exposure of the transcytosed pIgR-dimeric sIgA complexes thereby promoting retrograde recycling of the receptor [30–32, 34–36]and hence, a decrease in the transport of dimeric sIgA causing a diminished mucosal immune response to STEC O157 at the RAJ. This alternate method of immune suppression could allow ease of colonization and persistence by STEC O157 at the RAJ despite the pre-existing pIgR. All the aforementioned outcomes of a Slp-pIgR interaction require experimental validation, and studies are being planned for further exploration including the development of a Slp-targeting vaccine for use in cattle.

## MATERIALS AND METHODS

### Bacterial strains

The following strains from our previous study[26]were evaluated: **(i)** STEC O157 strain EDL932 (ATCC 43894: *stx* ^+^, *stx* ^+^, *eaeA*^+^, *hlyA*^+^, *slp^+^*; American Type Culture Collection/ATCC, Manassas, VA) referred to as EDL932-WT, **(ii)** STEC O157 Δ*slp* (*stx* ^+^, *stx* ^+^, *eaeA*^+^, *hlyA*^+^, *slp^-^*) referred to as EDL932 Δ*slp* and **(iii)** STEC O157 Δ*slp* + pUC18::*slp* (*stx* ^+^, *stx* ^+^, *eaeA*^+^, *hlyA*^+^, *slp^+^*, ampicillin resistant) referred to as EDL932 Δ*slp*-p:*slp*.

### Bacterial inoculum preparation

All bacteria were grown overnight in Dulbecco’s modified Eagle’s medium (DMEM) with low glucose (DMEM-LG; Invitrogen, Carlsbad, CA), with or without ampicillin (100 μg/ml), at 37°C without aeration. As described previously, the overnight cultures were washed and re-suspended in DMEM with no glucose (DMEM-NG; Invitrogen) prior to testing in the adherence assays [15, 43, 70, 71].

### RSE adherence assay

The RSE assay was done in three biological replicates with eight technical replicates per strain in each assay, as previously described [15–17, 19, 20, 43, 71, 83]. Briefly, RSE cells were suspended in DMEM-NG to a final concentration of 10^5^ cells/ml. Each bacterial isolate was mixed with RSE cells at a bacteria:cell ratio of 10:1. The mixture was incubated at 37°C with aeration (110 rpm) for 4 h, pelleted, washed, and reconstituted in 100 µl of double-distilled water (dH2O). Drops of the suspension (2 µl) were placed on Polysine slides (Thermo Scientific/Pierce, Rockford, IL), dried, fixed, and stained with fluorescence-tagged antibodies specific to the O157-antigen and cytokeratins of the RSE cells as before [15–17, 19, 20, 26, 71, 83]. Adherence patterns on RSE cells were qualitatively recorded as diffuse, aggregative, or nonadherent, and quantitatively as percentages of RSE cells with or without adhering bacteria [16]; a strongly adherent phenotype requires more than 50% of RSE cells with 10 adherent bacteria per cell, moderately adherent is when 50% or less of the RSE cells have 5 to 10 adherent bacteria per cell, and nonadherent when less than 50% of the RSE cells have only 1 to 5 adherent bacteria. RSE cells with no added bacteria were included as negative controls to confirm the absence of pre-existing STEC O157 bacteria. Quantitative data were evaluated for statistical significance using one-way ANOVA with Dunnett’s test and the unpaired t-test; p < 0.05 was considered significant (GraphPad Prism version 8.0.0, GraphPad Software, San Diego, CA).

### RAJ-IVOC adherence assay

Six RAJ-IVOC adherence assays were conducted, each in duplicate when sufficient tissue was available, with the test strains and a ‘no bacteria’ control. Bovine RAJ tissues for the IVOCs were collected at necropsies of animals in other studies at the National Animal Disease Center (NADC, Ames, IA) or from a local meat locker, with the approval of the NADC Institutional Animal Care and Use Committee (Supp. Table S1). Animals included Holstein, Angus, or Aberdeen cows or steers, 1-9 years of age, fed an alfalfa hay-based maintenance or corn-based finishing diet with ad libitum access to water (Supp. Table S1). The tissues were transported in DMEM-NG (Invitrogen) supplemented with 2.5% fetal bovine serum (FBS; Thermo Scientific HyClone, Logan, UT), 100 µg/ml streptomycin, 100 U/ml penicillin (Pen-Strep; Invitrogen), 2.5 mg/L amphotericin B (Sigma), and 50 µg/ml gentamicin (Invitrogen), on ice, and processed in the laboratory to either harvest RSE cells or to set up the RAJ-IVOCs as previously described [15, 43].

Briefly, the RAJ tissue was cut into multiple 4 cm x 2 cm (length x width) rectangular pieces, with the length encompassing 2 cm of each region on either side of the RAJ. The tissue pieces were carefully placed on top of a stack prepared in a flat-bottomed polystyrene clear tissue culture dish (Corning/Costar, Sigma-Aldrich Corp., St. Louis, Mo.) comprising of a sterile dental wax disc (Polysciences, Inc., Warrington, PA), a 1-2 mm thick sterile sponge soaked in DMEM-HG (Invitrogen) with 10% FBS and a sterile Whatman filter disc (Grade 1, 32 mm; Sigma) as shown in Supplementary Figure S1. The cut tissue piece was secured with sterile pins along the edges, with the mucosal/luminal surface facing outwards. Sterile DMEM-NG with 3% agarose was used to seal gaps around the edges of the tissue to prevent spillage of the DMEM-HG media below onto the exposed surface of the tissue and to contain the bacterial inoculum on the exposed luminal surface of the RAJ-IVOC (Supp. Fig. S1).

Given the estimated total of 10^4^ cells on the exposed surface area of the RAJ-IVOC, bacterial inoculum of 10^8^ CFU bacteria (10,000. bacteria:1 cell ratio) and 10^7^ CFU bacteria (1000 bacteria:1 cell ratio) in DMEM-NG (2 ml total volume) were evaluated in comparative assays to determine the optimal bacterial concentration producing distinct adherence phenotypes[43]. For no bacteria controls, 2 ml DMEM-NG without bacteria was added.

Post-inoculation, all RAJ-IVOC containing dishes were incubated at 39°C with 5% CO2 and gentle shaking at 100-110 rpm for 3 hours. Following incubation, the inoculum or media left on the exposed tissue surface was aspirated and plated on sorbitol MacConkey agar (BD Biosciences) containing 4-methylumbelliferyl-β-d-glucuronide (MUG, 100 mg/liter; Sigma) (SMAC-MUG) and MacConkey agar (BD Biosciences) containing MUG (100 mg/liter; Sigma) (MAC-MUG) to isolate STEC O157, and other background bacteria if any, respectively [43]. Each RAJ-IVOC tissue was then gently disengaged from the agarose, rinsed, and one half, weighing about 1-2 g, was frozen in 10 ml LB with 30% glycerol (LB-glycerol) for subsequent bacterial culture. The second half was flash frozen in Optimal Cutting Temperature solution (OCT; Tissue-Tek, Sakura Finetek, Torrance, CA) for sectioning and staining with hematoxylin and eosin (H&E) dyes at the Microscopy Services, NADC for histopathological evaluations, or with antibodies tagged with fluorescent tags to study the adherence of the inoculated bacteria [43]. Adherence patterns were qualitatively recorded as diffuse, aggregative (microcolonies), or nonadherent. The H&E-stained slides were analyzed using the Aperio Digital Pathology system (Leica Biosystems, Deer Park IL) and the corresponding Aperio Image Scope software (Leica).

### RAJ-IVOC tissue viability test

The viability of the RAJ-IVOC tissues was tested as described previously [43], using two uninoculated and unfixed tissue samples per assay, one pre-incubation, and the other post-incubation. The RedDot2 nuclear staining dye (RedDot™2 Far-Red Nuclear Stain, Biotium, Inc., Fremont, CA) was used per the manufacturer’s instructions. Briefly, each tissue sample was soaked in RedDot2 reagent (Biotium) diluted in DMEM-NG, for 30 min at room temperature, before rinsing and flash freezing in OCT. The stained frozen tissues sections, on Colorfrost slides (Thermo Fisher Scientific, Pittsburgh, PA) were air dried and cover slipped with Prolong Glass anti-fade reagent (Invitrogen) before visualization by fluorescent microscopy.

### RAJ-IVOC tissue culture for bacteria

The ∼1-2 g of RAJ-IVOC tissue frozen in LB-glycerol was thawed and cultured for STEC O157 as previously described [43, 84–86]. Briefly, the tissue was minced and suspended in 25-50 ml Trypticase soy broth (BD Bioscience, San Jose, CA), supplemented with cefixime (50 μg/L; U.S. Pharmacopeia. Washington, D.C.), potassium tellurite (2.5 mg/L; Sigma), and vancomycin (40 mg/L; Alfa Aesar, Haverhill, MA) (TSB-CTV), and the suspension was serially diluted with sterile saline (0.15 M NaCl) both before and after overnight incubation at 37°C with aeration. The pre-incubation dilutions were spread plated on SMAC-MUG (non-enrichment culture) and post-incubation suspension dilutions were plated on SMAC-MUG supplemented with cefixime (50 μg/L), potassium tellurite (2.5 mg/L) and vancomycin (40 mg/L) (SMAC-CTMV; selective-enrichment culture). All plates were read after overnight incubation at 37°C and colonies that did not ferment sorbitol or utilize MUG (non-fluorescent under UV light) were confirmed to be STEC O157 by latex agglutination tests (*E. coli* O157 latex, Oxoid Diagnostic Reagents, Oxoid Ltd., Hampshire, UK). The suspensions were additionally plated on MAC-MUG for increased recovery of the lactose-fermenting, MUG-utilizing (fluorescent under UV light) background non-STEC bacteria, if any. Bacteria recovered from the RAJ-IVOC tissue cultures were verified by PCR, as described below. Quantitative data from the comparative RAJ-IVOC tissue cultures were evaluated for statistical significance if any differences in adherence were observed using the unpaired t-test or one-way ANOVA with Dunnett’s test; p < 0.05 was considered significant (GraphPad Prism).

### PCR verification of strains.

Polymorphic amplified typing sequence (PATS) was used to DNA-fingerprint bacterial isolates, pre- and post-assays, as described previously [87–90] to confirm derivation from the STEC O157 strain EDL932. Specifically, primer pairs targeting 8 polymorphic *Xba*I-, 7 polymorphic *Avr*II- restriction enzyme sites, and 4 virulence genes encoding Shiga toxins 1 and 2 (*stx1* and *stx2*), intimin-γ (*eaeA*), and hemolysin-A (*hlyA*), were used to generate amplicons from colony lysates [20, 87, 88, 90, 91]. Purified (QIAquick PCR purification kit, Qiagen, Valencia, CA) PCR reactions amplifying the *Avr*II- restriction enzyme sites were digested with the *Avr*II restriction enzyme (New England Biolabs, Beverly, MA) to confirm the presence of the restriction site. Following 3% agarose gel-electrophoresis, the presence or absence of amplicons for *Xba*I sites and the virulence genes was recorded using “1” and “0”, respectively. For the *Avr*II site, absence of an amplicon was recorded as “0”, while the presence of the restriction site with a single nucleotide polymorphism was scored as “1”, an intact restriction site as “2”, and a restriction site duplication as “3” [87, 88, 91]. **(ii) Rapid differentiation** of the wild-type, mutant and complemented strains was done using primers targeting the vector pUC18, and the *slp* and *stx2* genes that were designed using the ‘PCR Primer Design’ tool in DNAStar (DNAStar Navigator Version 17.4.1.17; https://www.dnastar.com): **(i)** <u>pUC18 primers</u>: Forward 5′-TCGCGCGTTTCGGTGATGA-3′ and Reverse 5′-ACGAAAGGGCCTCGTGATACG-3′; amplicon size, 2685 bp, **(ii)** *<u>slp</u>* <u>primers</u>: Forward 5′-ATGAACATGACAAAAGGTGC-3′ and Reverse 5′-TTATTTGACCAGCTCAGGTGTTA-3′; amplicon size 567 bp, and **(iii)** *<u>stx2</u>* <u>primers</u> used for PATS profiling [87–90]: Forward 5′-GGCACTGTCTGAAACTGCTCC-3′ and Reverse 5′- TCGCCAGTTATCTGACATTCTG-3′; amplicon size 255 bp.

### Immunofluorescent staining of RAJ-IVOC tissue sections. (i) For test bacteria and host cells

As previously described [15, 43, 92], the tissue sections were ethanol-fixed, blocked with 5% normal goat serum, and incubated with primary and secondary antibodies, each at RT for 1 h, targeting the bacteria or host cell cytokeratin. Primary antibodies included the mouse anti-(PAN) cytokeratins (AbD Serotec, Raleigh, NC) targeting the RAJ cell cytokeratins. Secondary antibodies included the Alexa Fluor 594 (red)–labelled goat anti-mouse IgG (H+L; F(ab^’^)2fragment) (Invitrogen) targeting the anti-cytokeratins primary antibody and the fluorescein isothiocyanate (FITC; green)–labelled goat anti- O157 (KPL, Gaithersburg, MD) antibodies targeting O157. Air-dried slides were then cover-slipped with Prolong Gold anti-fade reagent containing the DNA stain 4^’^,6-diamidino-2-phenylindole (DAPI (blue); Invitrogen). Immunofluorescent images from the stained slides were captured using the Nikon Eclipse E800 fluorescence microscope (Nikon Instruments Inc., Elgin, IL) [15, 92]. Control slides with sections from uninoculated RAJ-IVOC were stained similarly to rule out nonspecific binding [15, 43, 92].

Additional slides were stained with only secondary antibodies to verify the specificity of the primary antibody and/or with FITC-tagged antibodies targeting unrelated *Salmonella* bacteria to demonstrate specificity of antibodies used, as needed. **(ii) For pIgR, EDL932-WT and host cells:** The tissue sections were fixed and blocked as before and stained with different combinations of primary and secondary antibodies. The rabbit anti-pIgR IgG (Thermo Scientific Pierce) targeting pIgR was the primary antibody.

Secondary antibodies included the Alexa Fluor 594 (red)–labelled goat anti-rabbit IgG (H+L; F(ab^’^)2fragment) (Invitrogen) targeting the anti-pIgR primary antibody and the fluorescein isothiocyanate (FITC; green)–labelled goat anti-O157 (KPL, Gaithersburg, MD) antibody targeting STEC O157. In addition, host cell cytokeratins were stained, when needed, with mouse anti-(PAN) cytokeratins (AbD Serotec, Raleigh, NC) primary antibody targeting the RAJ cell cytokeratins and the Alexa Fluor 488 (green) labelled goat anti-mouse IgG (H+L; F(ab^’^)2fragment) (Invitrogen) targeting the anti-cytokeratins primary antibody. Control slides were processed, and all stained slides were air-dried, cover-slipped and imaged as described above.

### RNAscope in situ hybridization (ISH) screening for pIgR-mRNA. (i) Tissue samples

The RNAscope ISH assay [93] was set up to screen the RAJ-IVOC tissues for the distribution and concentration of pIgR-mRNA. After a trial run with primarily the columnar epithelial region of the RAJ, RAJ-IVOC assays were set up with the following treatments: **#T0**: uninoculated, unincubated, pre-assay, **#1:** uninoculated, incubated, no bacteria control, or **#3:** with EDL932-WT, or **#4:** EDL932 Δ*slp*, or **#5**: EDL932 Δ*slp*-p:*slp* . All the RAJ-IVOC tissues were incubated for 3h, as described above, except for the #T0 IVOC which represented the pre-assay tissue. Each IVOC tissue sample, including #T0, was fixed overnight in neutral buffered formalin (NBF) before moving the formalin-fixed tissues into 70%ethanol. The formalin-fixed tissues were then embedded in paraffin for sectioning and use in the RNAscope ISH assay. **(ii)Assay.** RNAscope ISH assay was performed using the RNAscope® 2.5 HD Reagent Kit RED (Cat. No. 322750, Advanced Cell Diagnostics (ACD), Newark, CA) with the Leica Biosystems BOND RX automated IHC/ISH slide staining system (Leica), according to the manufacturer’s instructions. The formalin-fixed paraffin-embedded tissue sections (4–5 μm) were mounted on Superfrost Plus slides, baked at 60°C for 1 h, and subjected to automated deparaffinization with BOND Dewax Solution and graded alcohol rinses, followed by rehydration in BOND Wash Solution. Heat-induced epitope retrieval was carried out at 88°C using BOND ER Solution 2, after which sections were treated with RNAscope 2.5 LSx Protease and blocked with RNAscope 2.5 LSx H₂O₂. Hybridization was performed at 42 °C for ∼2 h with target-specific RNAscope™ 2.5 LS Probes including: (i) Test, Bt-pIgR-C1 (test; Cat No. 1268218-C1: 20 pairs complimentary to 354-1269 bp of the target pIgR mRNA-accession number NM_174143.1), (ii) Negative Control, dapB (Cat No. 312038: 10 pairs complimentary to 414-862 bp of the non-specific *Bacillus subtilis* strain SMY dapB mRNA-accession number, EF191515) and (iii) Positive control, Bt-PPIB (Cat No. 319458: 13 pairs complimentary to 24-788 bp of the host cell specific, *Bos taurus* cyclophilin B mRNA-accession number, NM_174152.2). Signal amplification was achieved through the sequential application of RNAscope AMP 1–6 RED reagents with intervening BOND Wash Solution rinses, and chromogenic detection was performed using the Mixed Red Refine reagent. Slides were counterstained with RNAscope Hematoxylin, blued with RNAscope Bluing reagent, rinsed in deionized water, and hydrated prior to coverslip mounting. All reagents were automatically dispensed in 150 µL volumes, and incubation times followed the ACD RNAscope 2.5 LSx standard protocol, as verified in the Leica BOND RX run setup log. **(iii) Digital image analysis.** Histologic sections were scanned at 400X magnification using an Aperio Versa 200 scanner. Digital image analysis was performed using HALO Image Analysis Platform version v3.6 and the *in-situ* hybridization module, ISH v3.4.3.0 (Indica Labs, Inc., Albuquerque, NM). To select the ROI for analysis, the RAJ was located followed by the selection of the SCE to one side and the SSE to the other side. The annotation was performed to ensure that both SCE and SSE had approximately the same length. For each sample, the ISH algorithm was first trained with representative nuclear and pIgR-ISH staining and then applied to the complete selected area. The following variables were obtained for SCE and SSE: total red (pIgR) copies, average red (pIgR) copies per cell, and percentage of pIgR positive cells. The ACD semi-quantitative scoring was also used to characterize expression levels, which ranges from 0 (no expression) to 4 (high expression, >15 dots/cell) (Supp. Table S2). The H-score reflects both the number of dots per cell and the percentage of cells displaying each specific dot count (Supp. Table S2).

### Protein modeling

The AlphaFold protein structure database (https://alphafold.ebi.ac.uk; [94, 95] and the Research Collaboratory for Structural Bioinformatics (RCSB) protein databank (https://www.rcsb.org; [96]) were used to access existing predicted structures or interactive models as needed. In addition, *de novo* protein-protein docking analysis was done using NovaDock in DNASTAR Protean 3D-Version 17.4.3 (2) (DNASTAR, Inc. Madison WI; https://www.dnastar.com). NovaDock employs the “SwarmDock” algorithm for computational modeling and makes docking predictions based on energy calculations and protein flexibility [46–48], generating 3D high-resolution models of the proposed receptor-ligand complexes. Protein structures for docking were selected from the AlphaFold database and included: (i) the ligand, *E. coli* Slp (UniProt: P37194; PDB ID: AF-P37194-F1-v4; DOI: https://alphafold.ebi.ac.uk/entry/P37194), (ii) the receptor, bovine pIgR (UniProt: P81265; PDB ID: AF-P81265-F1-v4; DOI: https://alphafold.ebi.ac.uk/entry/P81265), and (iii) the receptor, human pIgR (UniProt: P01833; PDB ID: AF-P01833-F1-v4; DOI: https://alphafold.ebi.ac.uk/entry/P01833) (Fig. 6; Supplementary PDB folder). The human pIgR-dimeric sIgA complex (PDB ID: 6UE7; DOI citation: https://doi.org/10.2210/pdb6UE7/pdb; [97]) was acquired from the RCSB protein database for comparison with the *de novo* docking models developed for human pIgR and *E. coli* Slp, and bovine pIgR and *E. coli* Slp proteins, using NovaDock. In addition, protein-protein alignments were done using BLASTp (https://blast.ncbi.nlm.nih.gov/) to verify sequence overlaps, if any.

## DATA AVAILABILITY

All relevant data has been included in the manuscript, either in the main or supplementary files.

## ACKNOWLEDGEMENTS

Excellent technical assistance provided by *Late* Mr. Bryan Wheeler, with tissue collection, experiments, data collection, and organization, is gratefully acknowledged. The support provided by NADC technicians and the Animal Resource Unit in providing access to RAJ tissue is acknowledged with appreciation. We sincerely acknowledge Adrienne Shircliff and Judith B. Stasko at the NADC Microscopy Services for expertly generating the H&E-stained image datasets in Aperio. We also thank Dr. Robert O. Ossiboff, Clinical Associate Professor at the College of Veterinary Medicine, University of Florida for lending his expertise in RNAScope ISH. This research was supported in part by the USDA-ARS CRIS project 5030-32000-225-00D (ITK). SK was supported by the College of Veterinary Medicine, University of Florida.

This research was supported by an appointment (ENB) to the Agricultural Research Service (ARS) Research Participation Program administered by the Oak Ridge Institute for Science and Education (ORISE) through an interagency agreement between the U.S. Department of Energy (DOE) and the U.S. Department of Agriculture (USDA). ORISE is managed by ORAU under DOE contract number DE- SC0014664. All opinions expressed in this paper are the authors’ and do not necessarily reflect the policies and views of USDA, ARS, DOE, or ORAU/ORISE. Mention of trade names or commercial products in this article is solely for the purpose of providing specific information and does not imply recommendation or endorsement by the U.S. Department of Agriculture. USDA is an equal opportunity provider and employer.

## SUPPORTING INFORMATION

### Supplementary Figure Legends

**Figure S1.**
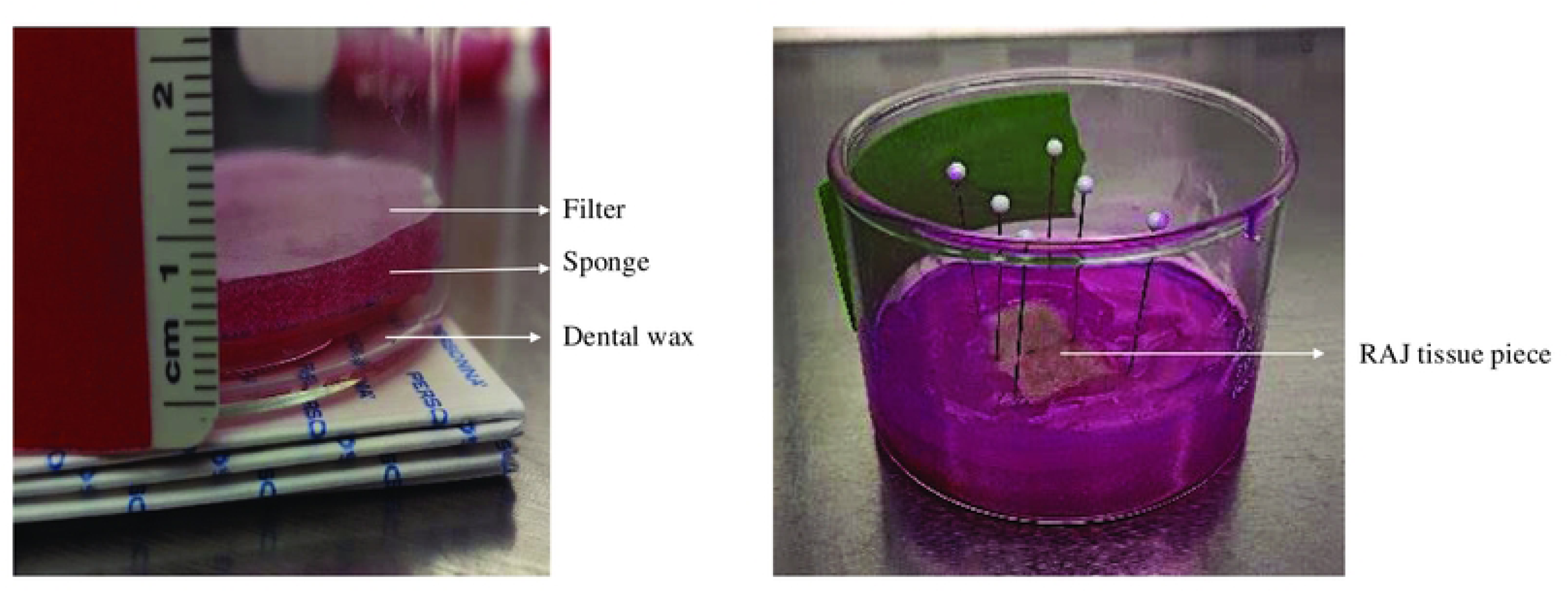
Assembled bovine RAJ- IVOC.

**Figure S2.**
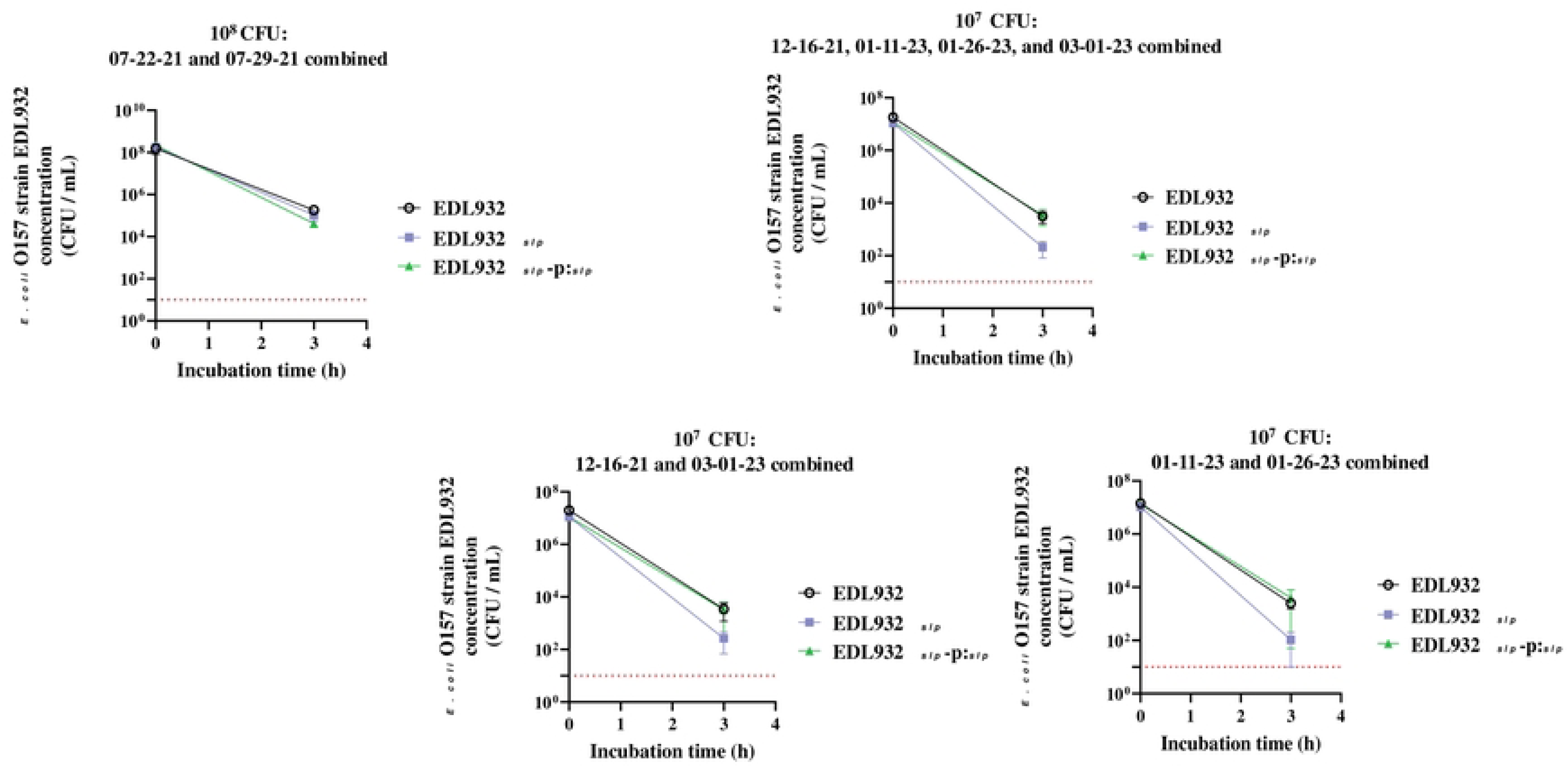
Graphs representing viable counts (CFU/ml) of test strains recovered from RAJ-IVOC tissues, by non-enrichment bacterial culture, from various assays with different inoculum concentrations. The culture counts were averaged between two assays at the 10^8^ CFU/ml inoculum and four assays at the 10^7^ CFU/ml inoculum. The red dotted line on the graph marks the STEC O157 detection limit of 10 CFU/ml for non-enrichment cultures.

**Figure S3.**
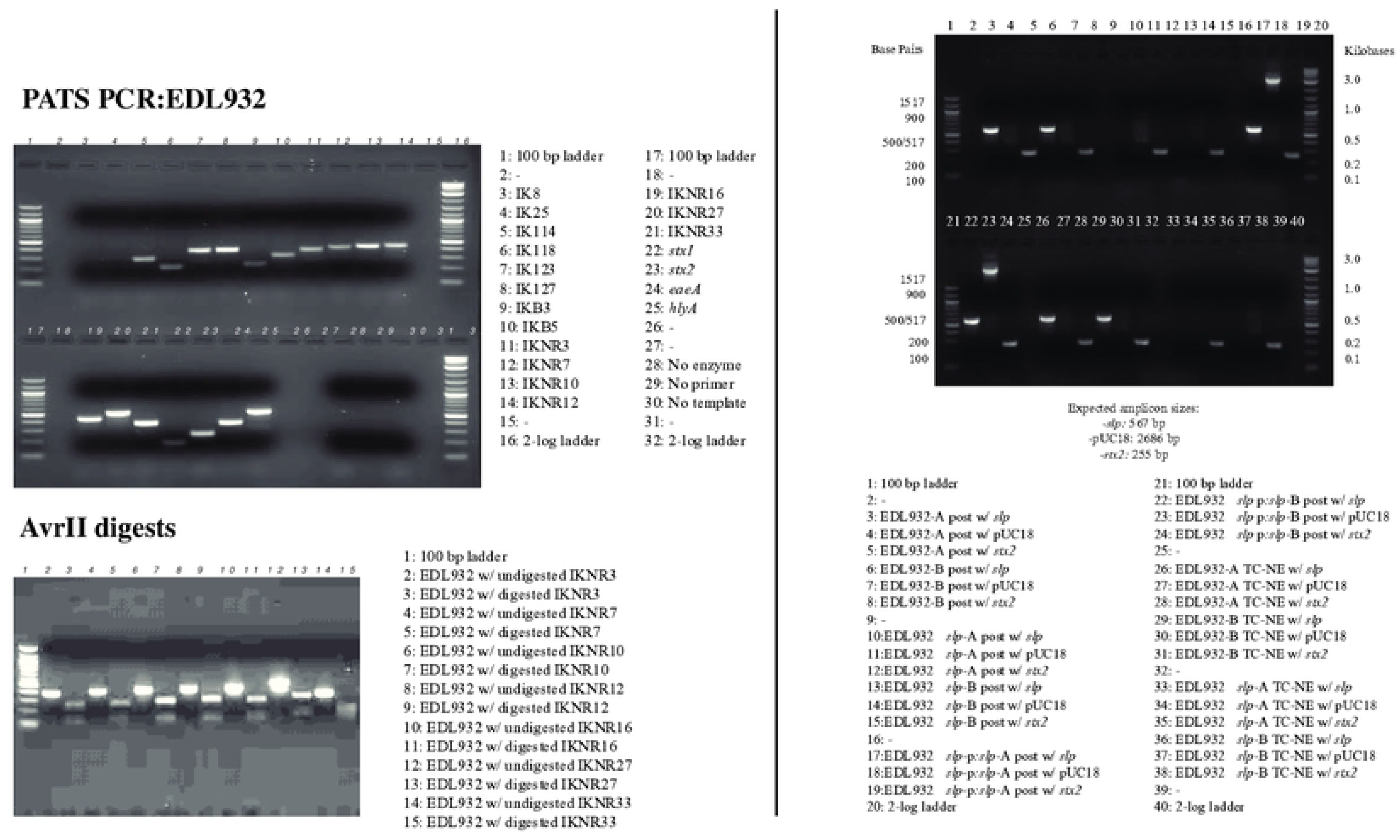
Electrophoretic patterns of representative PCR profiles on 3% agarose gels.

**Figure S4.**
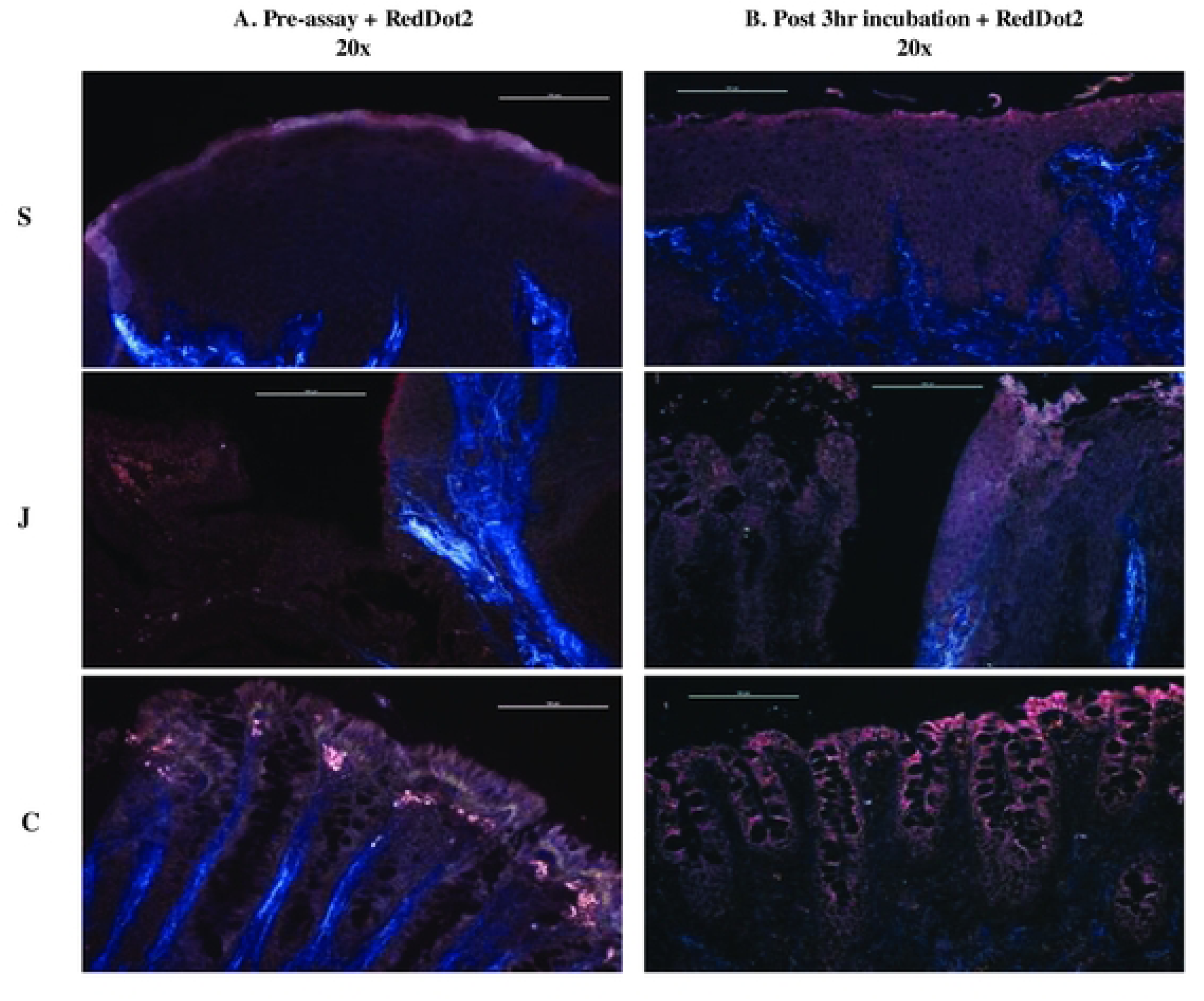
RAJ-IVOC viability test using the RedDot2 nuclei staining method. The un-fixed RAJ-IVOC tissue was stained with RedDot2 dye, (A) pre-assay and (B) post-3 h incubation. The squamous (S), junction (J), and columnar (C) regions of the stained RAJ-IVOC tissue are shown, along with the 100 µm scale bar. Images were captured at 200x magnification; the objective used is indicated on the images. The RedDot2 dye stains nuclei red in tissues with altered integrity. No red coloration of the nuclei, in both samples, reflects good tissue integrity and viability.

**Figure S5.**
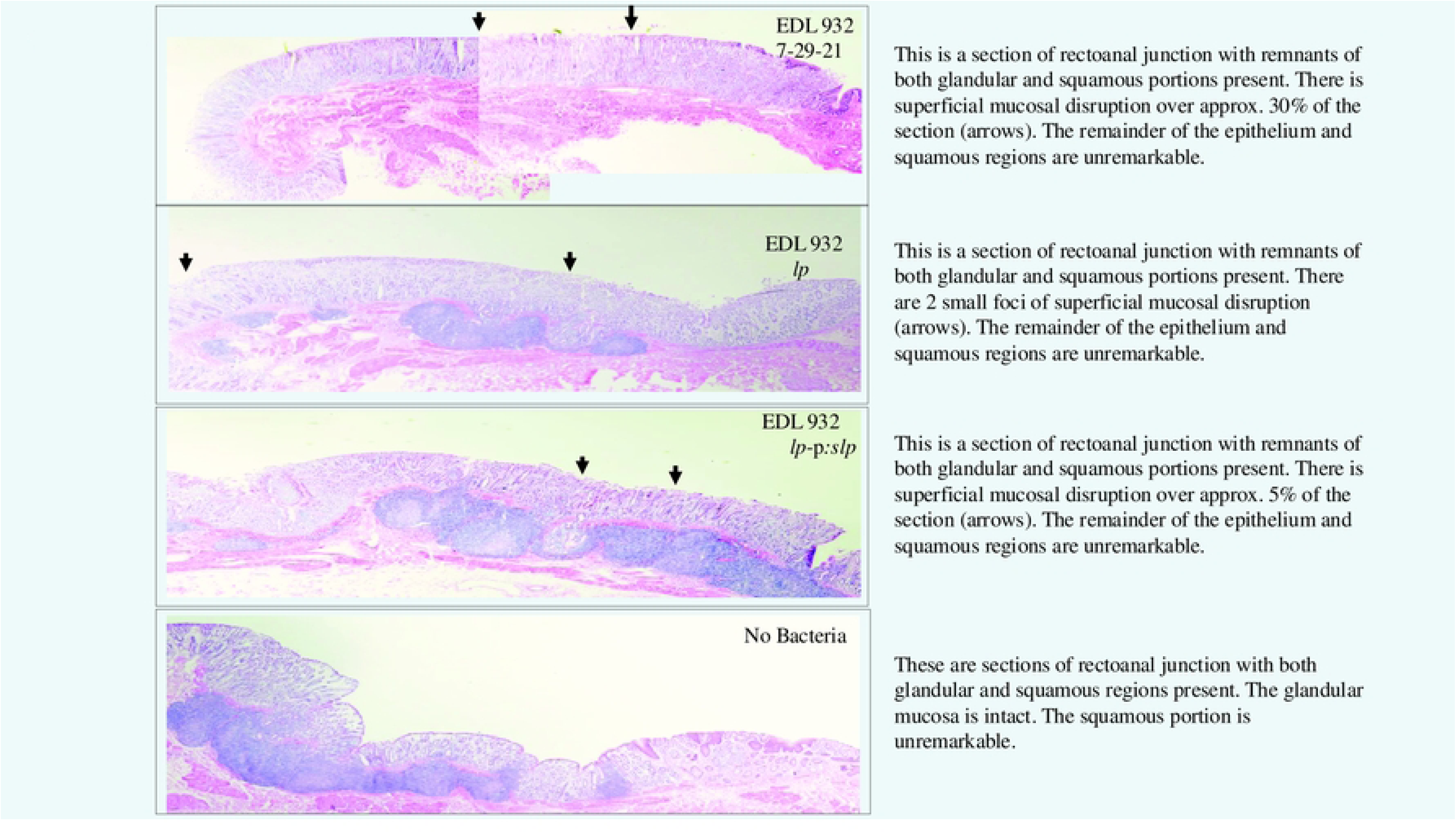
Representative histopathological report for a RAJ-IVOC assay. The RAJ-IVOC were inoculated with either EDL932-WT, EDL932 Δ*slp*, EDL932 Δ*slp*-p:*slp* or not inoculated (no bacteria), and incubated at 39°C for 3 h. H&E-stained tissue section slides were scanned using the Aperio digital pathology system to obtain the eImages. Additional detailed images and report are in the Supp. Histopathology Reports folder.

### Supplementary Tables

Supplementary Table S1. RAJ-IVOC assay data. Supplementary Table S2. Total RNAScope ISH data.

### Supplementary Data

1. Histopathology Reports.

2. RNAScope Images.

3. PDB files (Human pIgR-*E. coli* Slp and Bovine pIgR-*E. coli* Slp docking models).

Additional model and BLASTp information.

